# Supplier-origin gut microbiomes affect host body weight and select autism-related behaviors

**DOI:** 10.1101/2024.04.01.587648

**Authors:** Zachary L. McAdams, Kevin L. Gustafson, Amber L. Russell, Rachel Self, Amy L. Petry, Teresa E. Lever, Aaron C. Ericsson

**Affiliations:** Molecular Pathogenesis & Therapeutics Program, University of Missouri, Columbia, MO, 65201; MU Metagenomics Center, University of Missouri, Columbia, MO, 65201; Mutant Mouse Resource and Research Center at MU, Columbia, MO, 65201; Comparative Medicine Program, University of Missouri, Columbia, MO, 65201; Department of Veterinary Pathobiology, University of Missouri, Columbia, MO, 65201; Division of Animal Sciences, University of Missouri, Columbia, MO 65211; Department of Otolaryngology, School of Medicine, University of Missouri, Columbia, MO 65212; Department of Biomedical Sciences, University of Missouri, Columbia, MO 65211

**Author notes:** Corresponding Author: ACE.

**Keywords:** BTBR, ASD, Microbiome, Gut-Brain-Axis, Growth, Supplier-origin GM, Envigo, The Jackson Laboratory

## Abstract

Autism spectrum disorders (ASD) are complex human neurodiversities increasing in prevalence within the human population. In search of therapeutics to improve quality-of-life for ASD patients, the gut microbiome (GM) has become a promising target as a growing body of work supports roles for the complex community of microorganisms in influencing host behavior via the gut-brain-axis. However, whether naturally-occurring microbial diversity within the host GM affects these behaviors is often overlooked. Here we applied a model of population-level differences in the GM to a classic ASD model – the BTBR T^+^ Itpr3^tf^/J mouse – to assess how complex GMs affect host behavior. Leveraging the naturally occurring differences between supplier-origin GMs, our data demonstrate that differing, complex GMs selectively effect host ASD-related behavior – especially neonatal ultrasonic communication – and reveal a male-specific effect on behavior not typically observed in this strain. We then identified that the body weight of BTBR mice is influenced by the postnatal GM which was potentially mediated by microbiome-dependent effects on energy harvest in the gut. These data provide insight into how variability within the GM affects host behavior and growth, thereby emphasizing the need to incorporate naturally occurring diversity within the host GM as an experimental factor in biomedical research.

## Introduction

Autism spectrum disorders (ASD) are a collection of complex human neurodiversities characterized by reduced social communication and increased restrictive, repetitive behaviors^1,2^. The prevalence of ASD has risen to 1-in-36 children, with males being diagnosed at nearly four times the rate of females (4.3% vs 1.1%, respectively)^2^. In addition to the core ASD behaviors, many co-occurring conditions including anxiety, depression, and gastrointestinal disorders are frequently diagnosed in ASD patients^1,3^. The diversity of ASD behaviors and co-occurring conditions is attributed to the complicated and relatively unknown etiology of ASD as genetics, the environment, and interactions between the two factors influence both the incidence and severity of the neurodiversity^1^. Given the presence of gastrointestinal disorders in ASD patients and increasing evidence for microbiome-mediated effects on host behavior via the gut-brain-axis^4^, the gut microbiome (GM) has become a promising therapeutic target to improve quality-of-life for ASD patients.

The GM is the complex community of microorganisms colonizing the gastrointestinal tract, functioning in host metabolism, vitamin and short chain fatty acid production, and the synthesis of neuroactive compounds. Growing evidence supports crucial roles for the GM in modulating many host behaviors, including ASD-related behaviors^5,6^. For example, germ-free mice exhibit the core ASD-related behaviors (i.e., a lack of social preference and increased repetitive behaviors) which are reversed by repopulating the gut with complex microbial communities or even single, psychobiotic microbes (e.g., *Lactobacillus reuteri*)^7,8^. The use of ASD-specific mouse models also supports microbiome-mediated mechanisms in which microbial metabolites or modulation of host immune response affects host sociability, communication, and stereotypic behavior^5,9,10^. While much of this work has focused on therapeutic roles for individual psychobiotic microorganisms or microbial metabolites, few have acknowledged how complex, naturally occurring differences in the composition of the GM influence ASD-related behaviors.

The naturally-occurring differences in GM communities between commercial rodent producers can be leveraged as a model of population-level differences in microbiome. The large differences in microbial diversity between GMs originating from The Jackson Laboratory and Envigo, in particular, are associated with robust effects on multiple host phenotypes in CD-1 mice including anxiety-related behavior, feeding behaviors, *in utero* growth, and adult body weight^11–13^. In addition to behavior and growth, supplier-origin GMs have also been found to differentially affect host immunity and disease susceptibility^14–16^. Interestingly, other groups have found that comparable supplier-origin communities affect the ASD-related behaviors of the maternal immune activation (MIA) mouse model of ASD^9,17^. Maternal T helper type 17 (Th17) immune responses in pregnant mice with a Taconic-, but not Jackson Laboratory-origin microbiome produce offspring exhibiting greater ASD-related behaviors^9^. The application of these distinct supplier-origin communities in the MIA model of ASD provided a unique platform to identify a single bacterial taxon sufficient to induce maternal Th17 responses and ASD-related behavior in the offspring^9,18^.

Here we utilized a similar discovery-based microbiome model to determine whether supplier-origin GMs originating from The Jackson Laboratory or Envigo affect the ASD-related behavior and growth of the BTBR T^+^ *Itpr3^tf^*/J (BTBR) mouse. The BTBR mouse exhibits robust ASD-related behaviors including altered ultrasonic vocalizations (USVs), increased repetitive behaviors, and a lack of social preference^19–23^. Interestingly, this model also demonstrates altered gut physiology and GM composition, thus increasing its utility in investigating the role of the GM in ASD-related behaviors^24,25^. Our approach leveraged BTBR mice colonized with The Jackson Laboratory- and Envigo-origin GMs. Using a robust panel of neonatal and adult ASD-related behavioral testing approaches, we identified selective supplier-origin GM-dependent effects on host ASD-related behavior – especially neonatal ultrasonic communication. We also found that the body weight of BTBR mice is influenced by the postnatal GM and that this effect may be mediated by microbiome-dependent effects on feed conversion in the gut.

## Results

### Supplier-origin microbiomes selectively affect ASD-related behavior

We characterized the ASD-related behavior of BTBR mice colonized with two supplier-origin GMs (**Figure 1A**). Relatively speaking, the GM originating from The Jackson Laboratory was less rich than the GM representative of Envigo (*p*_GM_ < 0.001, **Figure 1B**), thus, these communities were referred to as GM_Low_ and GM_High_, respectively. Sex-dependent effects on richness were also observed (*p*_Sex_ = 0.002). While these communities did not differ in alpha diversity (*p*_GM_ = 0.867, **Figure 1C**), significant differences in both beta diversity (*p*_GM_ < 0.001, **Figure 1D**) and taxonomic composition (**Figure S1**) were observed. Using ALDEX2^26^, sixty-one genera (37%) were identified as differentially abundant between the two communities, including an uncultured *Peptococcaceae* genus and *Anaeroplasma* from the phylum *Bacillota* enriched in GM_Low_ and *Mucispirillum* (phylum *Deferribacterota*) and *Bilophila* (phylum *Desulfobacterota*) enriched in GM_High_ (**Supplementary File 1**). Differentially abundant taxa were confirmed using ANCOM-BC2^27^ (**Supplementary File 1**).

**Figure 1.**
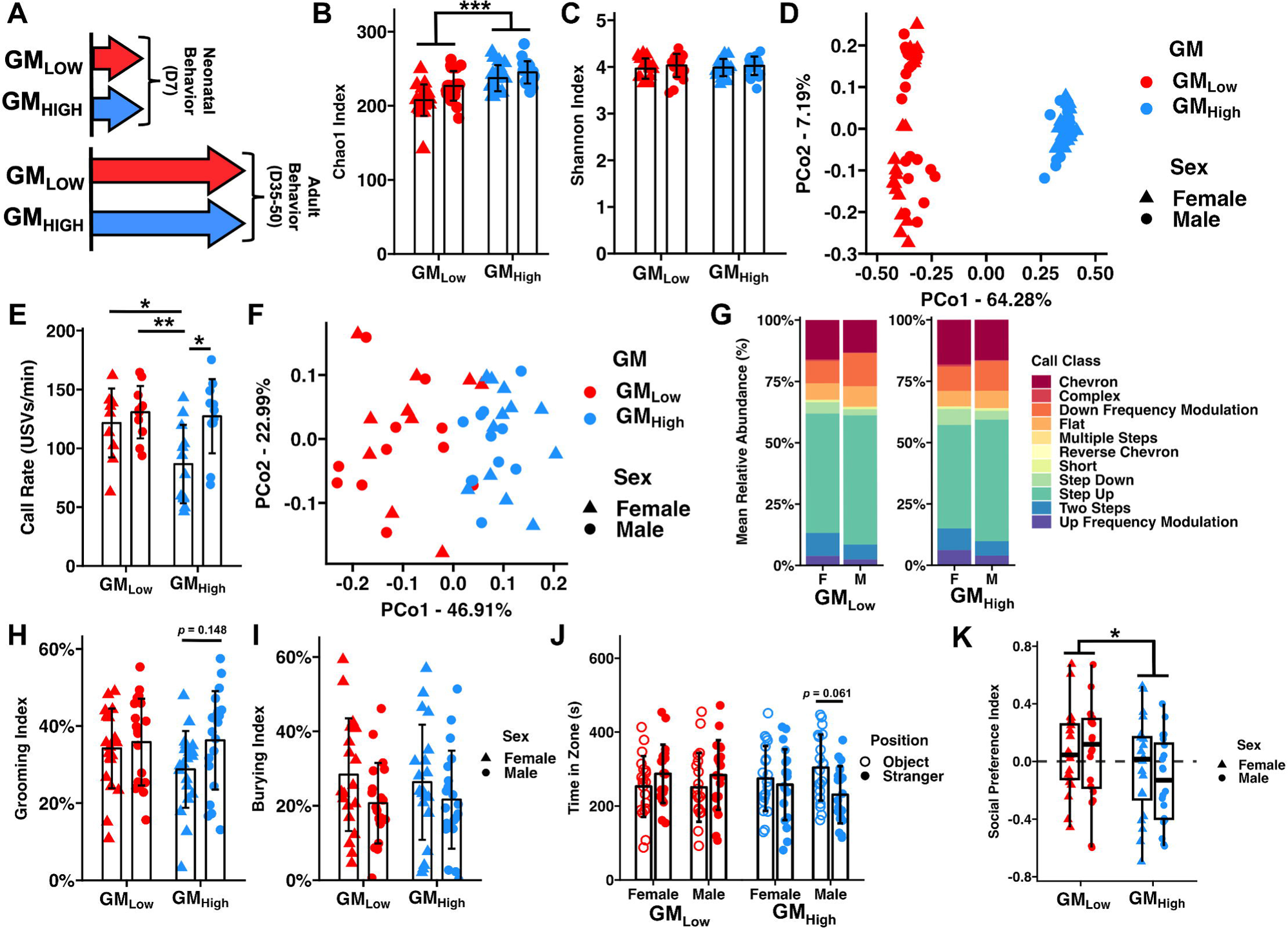
Standardized complex microbiomes selectively affect male ASD-related behavior. (A) Graphical representation of experimental design depicting cohorts of neonatal (*n* = 10-12 mice/sex/GM) and adult (*n* = 20 mice/sex/GM) BTBR mice. (B) Dot plot depicting Chao-1 Index. *** *p*_GM_ < 0.001, *p*_Sex_ = 0.002, Two-way ANOVA (C) Dot plot depicting Shannon Index. *p*_GM_ *=* 0.867, *p*_Sex_ = 0.276, Two-way ANVOA (D) Principal coordinate analysis depicting Bray-Curtis dissimilarity between microbial communities. *p*_GM_ < 0.001, *p*_Sex_ = 0.063, Two-way PERMANOVA. (E) Dot plot depicting USV rate. * *p* < 0.05, ** *p* < 0.01, Tukey *post hoc*. (F) Principal coordinate analysis depicting Bray-Curtis dissimilarity of the relative abundance of ultrasonic vocalizations. *p*_GM_ = 0.001, *p*_Sex_ = 0.044, Two-way PERMANOVA. (G) Stacked bar charts depicting mean relative abundance of call types determined by VocalMat. (H) Dot plot depicting Grooming Index. *p*_GM_ = 0.321, *p*_Sex_ = 0.069, Two-way ANVOA. (I) Dot plot depicting Burying Index. *p*_GM_ = 0.862 *p_Sex_* = 0.048, Two-way ANOVA. (J) Dot plot depicting time spent in Stranger (closed circles) or Object (open circles) chambers of social preference test. (K) Tukey box plot depicting Social Preference Index. * *p*_GM_ *=* 0.044, *p*_Sex_ = 0.498, Two-way ANOVA.

We first assessed ultrasonic vocalizations (USVs) in neonatal BTBR mice (n = 10-12 mice/sex/GM). A significant, albeit subtle, GM-dependent effect (*p*_GM_ = 0.038) on USV rate was observed with GM_Low_ BTBR mice exhibiting a greater USV rate than GM_High_ mice (**Figure 1E**). While a significant sex-dependent effect (*p*_Sex_ = 0.006) on USV rate was also observed, Tukey *post-hoc* testing revealed an interesting interaction of sex and GM within GM_High_ mice where males exhibited a greater USV rate than females (*p* = 0.011), suggesting greater ASD-related behavior in this group. This sex-dependent difference was not observed in GM_Low_ mice. Significant GM-(*p*_GM_ < 0.001) and sex-dependent effects (*p*_Sex_ = 0.044) on the overall USV repertoire were observed (**Figure 1F**). Specifically, significant sex-dependent effects on the relative abundance of “complex” and “step down” calls were observed, whereas the relative abundance of “up frequency modulation” calls differed by both sex and GM (**Figure 1G**, **Supplementary File 2**).

In separate cohorts of adult BTBR mice (*n* = 20/sex/GM) we then assessed repetitive and social behaviors. Using the self-grooming test (**Figure 1H**), we observed no GM-dependent effects (*p*_GM_ = 0.321) on grooming behavior; however, a strong sex-dependent trend (*p*_Sex_ = 0.069) was observed, with males exhibiting greater grooming behavior than females. Conversely, female mice exhibited significantly greater (*p*_Sex_ = 0.048) burying behavior than males (**Figure 1I**). No GM-dependent effects of burying behavior were observed (*p*_GM_ = 0.862). Lastly, we assessed social behavior using the three-chamber social preference test. As expected of the BTBR model of ASD^20^, we observed no overall differences in time spent between the stranger and object chambers (*p*_Position_ = 0.685). Additionally, neither GM (*p*_GM_ = 0.892) nor sex (*p*_Sex_ = 0.951) affected time spent in either chamber of the social preference test overall; however, a *post hoc* analysis using paired T tests revealed a strong trend towards GM_High_ males exhibiting greater asocial behavior (*p* = 0.061), spending more time in the object zone relative to the stranger zone. Finally, we determined the social preference index (SPI = [time_stranger_ - time_object_] / [time_stranger_ + time_object_])^28^ and found that BTBR mice with GM_High_ exhibited significantly reduced sociability compared to mice with GM_Low_ (*p*_GM_ = 0.044, **Figure 1K**). Collectively, these data suggest that standardized complex GMs selectively affect ASD-related behaviors of the BTBR mouse.

GM-, age-, and sex-matched C57BL/6J (B6) mice used as behavioral controls in our adult ASD-related behavior testing also exhibited select GM-dependent effects on ASD-related behavior. No GM-dependent effects on B6 grooming behavior were observed; however, GM_High_ B6 mice exhibited significantly reduced burying activity relative to GM_Low_ B6 mice (**Figure S2A-B**). B6 mice overall exhibited the social behavior expected of the strain, spending more time in the stranger zone compared to the object zone; however, no GM-dependent effects on social behavior were observed (**Figure S2C-D**).

### Standardized complex GMs postnatally affect body weight

While assessing the effect of supplier-origin GMs on the ASD-related behavior of BTBR mice, we collected body weights as previous work by our group using comparable GMs in CD-1 mice has revealed microbiome-dependent effects on body weight^12,13^. In the cohort of neonatal mice used for USV testing, we measured body weight at birth (D0) and after testing (D7). A total of 10 litters were weighed (5 GM_Low_ and 5 GM_High_) at birth. Litters ranged from 5 to 12 pups (9.5 ± 2.3) with no GM-dependent effects on litter size (*p*_GM_ = 0.383, T test). Following the collection of birth weights, litters were culled to 8 mice with an equal representation of males and females when possible.

At birth, GM_High_ BTBR mice were significantly heavier than GM_Low_ mice (*p*_GM_ < 0.001, **Figure 2A**). A similar GM-dependent effect on body weight was observed at D7 where GM_High_ mice were again significantly heavier than GM_Low_ BTBR mice (*p*_GM_ = 0.004). In the cohorts of BTBR mice used for adult behavior testing, however, we observed that GM_Low_ mice were heavier at weaning (D21, *p*_GM_ < 0.001, **Figure 2C**) and adulthood (D50, *p*_GM_ = 0.078, **Figure 2D**). Interestingly, B6 mice colonized with GM_Low_ also weighed more than those with GM_High_ at weaning (*p*_GM_ < 0.001, **Figure S3A**) and adulthood (*p*_GM_ = 0.069, **Figure S3B**).

**Figure 2.**
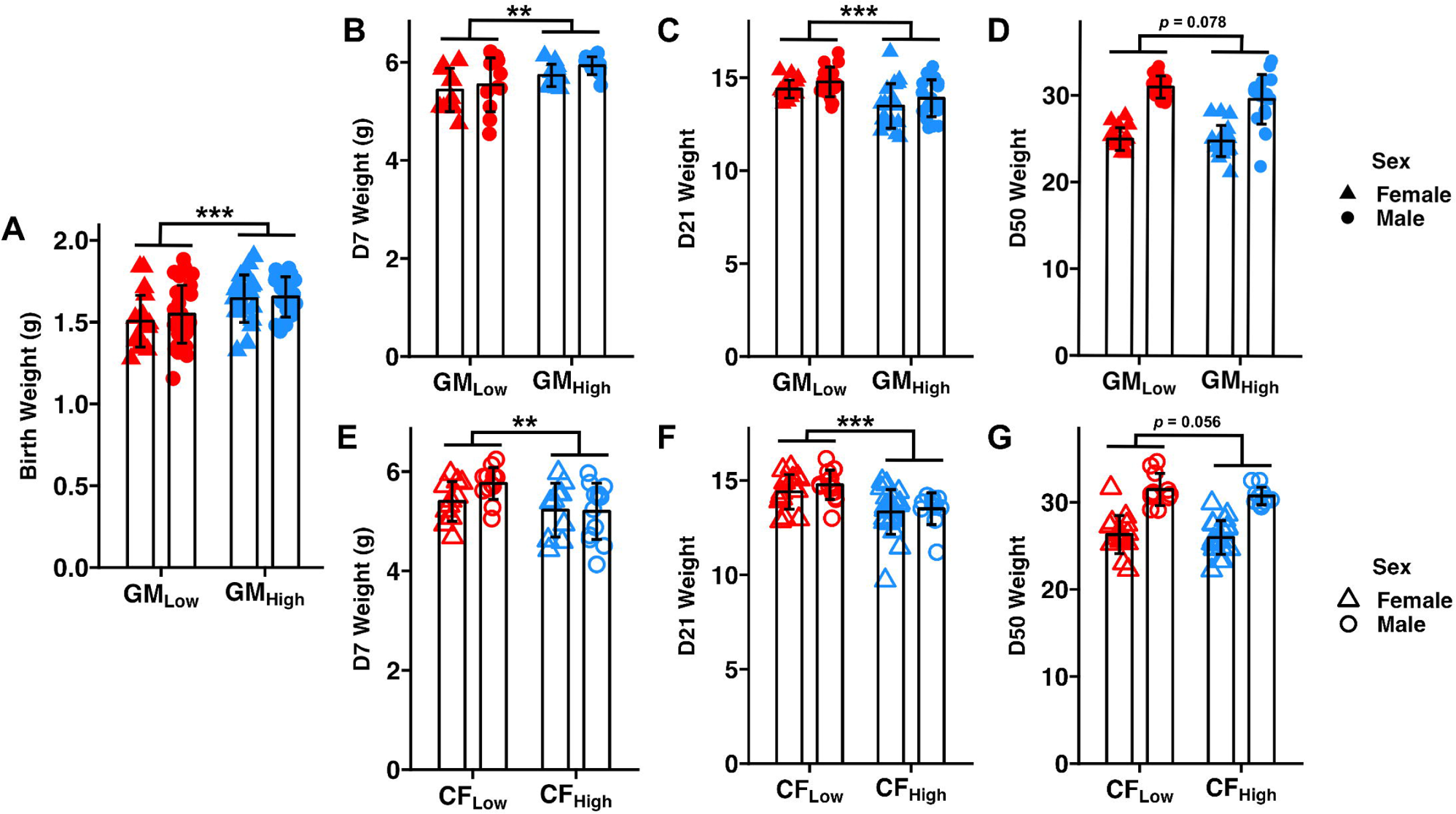

Given that BTBR mice born to a GM_Low_ dam weighed less than pups born to a GM_High_ dam at birth but were heavier in adulthood, we hypothesized that the postnatal GM influenced body weight. To confirm that the postnatal microbiome influenced body weight, we employed a cross-fostering experimental approach wherein mice born to GM_Low_ or GM_High_ dams were cross-fostered onto surrogate dams of the opposite GM within 48 hours of birth (**Figure S4A**)^29^. Mice born to a GM_Low_ birth dam but cross-fostered to and raised on a GM_High_ surrogate dam were referred to as CF_High_ (meaning “cross-fostered” onto GM_High_) with the reciprocal group (i.e., born to GM_High_ but cross-fostered onto GM_Low_) being referred to as CF_Low_ (meaning “cross-fostered” onto GM_Low_). If the observed GM-dependent effect on body weight was influenced by the prenatal (i.e., maternal) GM, then the phenotype would match that of the adult birth dam. Conversely, if this phenotype is influenced postnatally, then the phenotype would match that of the adult surrogate dam.

We confirmed that cross-fostering successfully transferred the GM from surrogate dam to cross-fostered mice using 16S rRNA sequencing of fecal samples collected at fifty days of age. Cross-fostered mice exhibited similar taxonomic composition to the surrogate dams (**Figure S5**). Additionally, alpha and beta diversity of these mice were characteristic of the fostered microbial communities, with CF_Low_ mice exhibiting a less rich and compositionally distinct GM compared to CF_High_ mice (**Figure S4B-D**). In cross-fostered BTBR mice, animals with CF_Low_ were heavier than CF_High_ at PND7 (*p*_GM_ = 0.006, **Figure 2E**). In separate cross-fostered cohorts, however, CF_Low_ mice were heavier at weaning (*p*_GM_ < 0.001, **Figure 2F**) and adulthood (*p*_GM_ = 0.056, **Figure 2G**). Given that the GM-dependent effect on body weight was similar to the phenotype of the mature surrogate dam GM, these data support that these supplier-origin GMs postnatally affect body weight in BTBR mice.

### Cross-fostering abrogates select effects on BTBR ASD-related behavior

Previous reports have shown that the maternal *in utero* BTBR environment contributes to offspring ASD-related behavior of the model^30,31^, thus we sought to determine whether the selective GM-dependent effects on ASD-related behavior (**Figure 1E-K**) were programmed *in utero* by the maternal GM or influenced, like body weight, primarily by the postnatal GM. In neonatal mice (n = 10-13 mice/sex/GM), no significant differences in USV call rate or composition were observed between groups (**Figure S4E-F**); however, GM-dependent effects on the relative abundance of “step up”, “step down”, and “up frequency modulation” calls were observed (**Figure S4G, Supplementary File 3**). While adult (11-21 mice/sex/GM) grooming behavior was not affected by GM (*p*_GM_ = 0.237, **Figure S4H**), CF_Low_ mice exhibited greater repetitive burying activity than CF_High_ BTBR mice (*p*_GM_ = 0.049, **Figure S4I**). Social behaviors did not differ between CF_Low_ and CF_High_ BTBR mice (**Figure S4J**). Collectively, the select GM-dependent effects of ASD-related behavior of BTBR mice were abrogated by cross-fostering, suggesting the ASD-related behaviors of BTBR mice may be influenced by factors from both the pre-(i.e., maternal) and postnatal GM.

### Supplier-origin GMs potentially affect nutrient acquisition in the BTBR mouse

To explore potential mechanisms influencing the postnatal GM-dependent effect of body weight of BTBR mice, we assessed three facets of host energy balance: food intake, voluntary activity, and fecal energy loss. We first assessed food intake by measuring the relative food intake of standard maintenance chow (LabDiet #5053 Chow) in pair-housed mice (*n* = 12-16 mice/sex/GM, *n* = 6-8 cages/sex/GM) for six weeks, beginning at weaning. As expected, BTBR mice with GM_Low_ weighed more than those with GM_High_ throughout the food intake experiment (*p* < 0.001, **Figure S6A**). Despite the difference in body weight, these groups consumed similar amounts of food over the six-week period (**Figure S6B**). Given that food intake positively correlated with body weight in both GMs (GM_Low_ ρ = 0.67, *p* < 0.001; GM_High_ ρ = 0.49, *p* < 0.001; **Figure S6C**), we determined feed efficiency by normalizing food intake at the cage level by the combined body weight of mice in the cage; however, no GM-dependent effects on feed efficiency were observed (**Figure 3A**). Interestingly, female mice exhibited a higher feed efficiency than males (*p*_Sex_ < 0.001).

**Figure 3.**
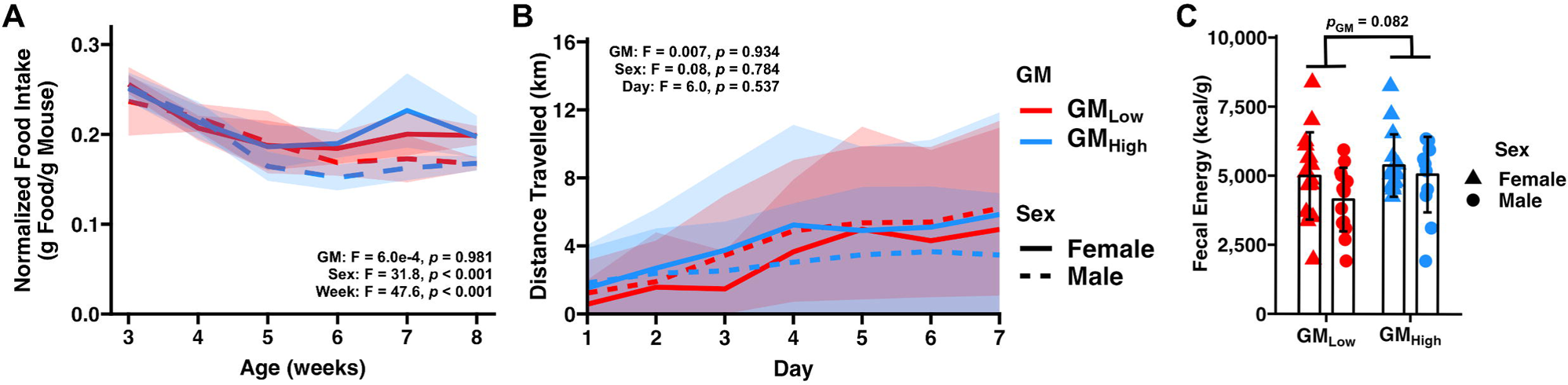
Standardized complex microbiomes may affect energy harvest in the gut. (A) Line plot depicting feed efficiency observed in GM_Low_ and GM_High_ BTBR mice. Bold line represents average feed efficiency. Ribbon represents standard deviation. Inset depicts three-way ANOVA results. (B) Line plots depicting distance traveled by GM_Low_ and GM_High_ BTBR mice. Bold line represents average distance travelled. Ribbon represents standard deviation. Inset depicts three-way ANOVA results. (C) Dot plot depicting time fecal energy as determined by bomb calorimetry. *p*_GM_ = 0.082, *p_Sex_* = 0.093, Two-way ANOVA.

Turning to mechanisms of energy loss, we measured output using voluntary running wheel activity of individually housed mice for one week. No sex- or GM-dependent effects on total distance travelled were observed, suggesting no difference in physical activity levels between GM_Low_ and GM_High_ BTBR mice (**Figure 3B**). Lastly, we measured fecal energy loss using bomb calorimetry of fecal samples collected over the course of a two- to three-day period. While no significant differences were observed, strong sex-(*p* = 0.093) and GM-dependent (*p* = 0.082) trends on fecal energy were observed. GM_High_ mice exhibited greater fecal energy content indicating reduced energy harvest in the gut. Collectively, these data suggest that effects of supplier-origin GMs on energy harvest from the diet may contribute to the postnatal GM-mediated effect on body weight.

## Discussion

Our data demonstrate selective GM-dependent effects on the behavior and growth of the BTBR mouse model of ASD. Specifically, we observed GM-dependent effects on vocalization rate and call composition in neonatal mice and multiple strong trends in adults that collectively suggest an Envigo-origin microbiome (GM_High_) exacerbated the ASD-related behavior of male BTBR mice. While the overall effects on behavior were selective and largely subtle, we revealed that the mature postnatal GM influenced body weight, beginning at weaning and persisting into adulthood. Mature BTBR mice with a low richness, Jackson Laboratory-origin microbiome weighed more than those with an Envigo-origin GM. While no GM-dependent effects on food intake or voluntary activity were observed, our data indicate that the postnatal GM may affect body weight by modulating energy acquisition in the gut. Collectively, these data suggest that the BTBR mouse model of ASD is susceptible to GM-mediated effects on both behavior and metabolism.

Separation- or stress-induced USVs have long been used as a measure of neonatal ASD-related behavior; however, the translatability of this phenotype is difficult to interpret as both the vocalization rate and call repertoire are highly variable across mouse strains and models of ASD^21,32^. For example, vocalization rate is often increased in the BTBR and MIA ASD models but decreased in some transgenic models of ASD (e.g., *Cntnap2^−/−^*), yet both effects on vocalization rate are classically defined as an ASD-related behavior associated with communication^9,21,33^. While the etiology of this behavior is unclear, our data demonstrate that variability in the literature regarding the USV phenotype may be influenced, in part, by the host GM. Whether these effects are due to microbiome-mediated influence on the neonatal stress response or even maternal care remains unknown; however, our data support that the communication ASD-related phenotype measured in neonatal BTBR mice is influenced by the GM.

Despite ASD being diagnosed more frequently in male patients, a similar sex-bias is not consistently observed across mouse models of the neurodiversity. Historically, the BTBR mouse presents strong ASD-related behaviors in both males and females, indicating the strain may not be a useful model of the ASD sex bias. Our behavior data suggest that the GM preferentially exacerbates male ASD-related behavior and that the typical Jackson Laboratory-origin GM may contribute to the lack of sex-dependent differences in the presentation of ASD-related behaviors. In the present study, BTBR mice with a Jackson Laboratory-origin GM (GM_Low_) exhibited no sex-dependent differences in ultrasonic communication, repetitive, or social behaviors; however, when colonized with an Envigo-origin GM (GM_High_), BTBR mice demonstrated male-specific increases in all three of the core ASD-related behaviors. Multiple genetic and hormonal mechanisms have been proposed to explain the strong male bias in ASD diagnoses^34–36^; however, the contribution of the GM to this sex bias in mouse models of ASD, let alone humans, is yet to be described.

The postnatal GM-mediated effects on body weight identified in this study were of particular interest. In CD-1 mice, we have historically found that animals with a Jackson-origin microbiome (GM_Low_) are heavier than those with an Envigo-origin (GM_High_), beginning *in utero* and persisting into adulthood^12,13^. The GM-dependent difference in body weight observed in CD-1 mice is likely due to an effect on overall growth, as these groups do not differ in relative body composition and GM_Low_ CD-1 mice exhibit increased cardiac weight^12^. Rather than an *in utero* programming of body weight – as in CD-1 mice – we found that the body weight of BTBR mice is postnatally influenced by the GM, as GM_High_ BTBR mice were heavier at birth but weighed less than GM_Low_ animals in adulthood (**Figure 2**). Fetal growth and neurodevelopment are influenced, in part, by the maternal microbiome modulating placental vascularization and nutrient availability to the fetus^37,38^, and given that the maternal BTBR *in utero* environment contributes to the development of ASD-related behavior^31^, it is reasonable to hypothesize that the maternal GM of BTBR mice may influence both the fetal growth and development of ASD-related behaviors in BTBR offspring.

Adult body weights exhibited the expected phenotype of their respective GM; mature BTBR mice with GM_Low_ were heavier those with GM_High_. Further supporting these GM-dependent effects on adult body weight, we observed similar GM-dependent effects in age- and sex-matched B6 mice (**Figure S3**). Whether the GM-dependent effect on body weight in either strain is due to differences in body size or composition remains unknown; however, it may be strain-specific as the BTBR mouse exhibits increased abdominal obesity and peripheral insulin resistance relative to B6 mice, both of which are factors that may be influenced by the host GM^39–41^.

Contrary to our conclusion that the postnatal GM influences body weight, we observed that – consistent with the body weight phenotype at birth – BTBR mice born to GM_High_ dams were heavier than those born to GM_Low_ dams at one week of age. This is likely due to the GM having not yet matured to the point at which the postnatal GM could influence body weight. The gastrointestinal tract of neonatal mice undergoes tremendous development during the first few weeks of life as the gut transitions from a highly aerobic to anaerobic environment and the host moves from maternal sources of nutrition to solid food^42,43^. Consistent with this idea, the neonatal GM at seven days of age is more similar to that of the oral microbiome of the dam than the fecal microbiome, including several aerobic bacterial taxa like *Lactobacillus* and *Streptococcus* dominating the neonatal gastrointestinal tract of CD-1 mice^44^. The pup fecal microbiome does, however, become more similar in composition to the maternal fecal microbiome around three weeks of age^44^, the same age at which we observed postnatal GM-dependent effects on body weight in BTBR mice. Collectively, our body weight data suggest that the mature (post-weaning) postnatal GM affects the body weight of the BTBR mouse.

When identifying the mechanism by which the postnatal GM influenced body weight in BTBR mice, we observed no effects on food intake or voluntary activity; however, a strong trend towards a GM-dependent effect on energy harvest was identified (**Figure 3**). GM_High_ BTBR mice excreted more fecal energy than GM_Low_ mice, which is consistent with the hypothesis that this group extracted fewer calories from the diet, leading to the decreased weight relative to GM_Low_ mice. Acknowledging that both the host and microbiome harvest energy from the diet^45,46^, multiple microbiome-mediated mechanisms may be working in concert to influence host body weight, including the alteration of host gene expression within the gut modulating nutrient availability and absorption^47,48^. Alternatively, the diverse members of these bacterial communities may have differing energy requirements for replication which may, in turn, affect energy availability to the host. Exploring these mechanisms may reveal novel microbiome-mediated mechanisms influencing feed conversion with profound metabolic and economic implications.

Collectively, our data have implications regarding both the specific use of the BTBR mouse in biomedical research and more broadly to behavioral and metabolic research involving the GM. Specific to the BTBR mouse, we have demonstrated that this model of ASD is susceptible to selective GM-dependent effects on the core ASD-related behaviors (particularly in males) and that body weight is influenced by the postnatal GM. The exact microbiome-mediated mechanisms driving these differences in behavior and growth in this model are yet to be determined, but complex microbial communities should be considered when using the BTBR mouse. More broadly, this work emphasizes the need to incorporate complex communities into gut-brain-axis research as it adds to a growing body of literature demonstrating that variability within the host GM contributes to variability of host phenotypes.

## Methods

### ETHICS STATEMENT

This study was conducted in accordance with the recommendations set forth by the Guide for the Care and Use of Laboratory Animals and was approved by the University of Missouri Institutional Animal Care and Use Committee (MU IACUC protocol 36781).

### ANIMALS

BTBR (RRID:IMSR_JAX:002282) and C57BL/6J (RRID:IMSR_JAX:000664) mice were purchased from The Jackson Laboratory (Bar Harbor, ME, USA). Mice were bred and pups were cross-fostered onto CD-1 surrogate dams within 24 hours of birth. The CD-1 surrogate dams were acquired from a colony of mice colonized with an Envigo-origin microbiome (GM_High_) maintained at the NIH-funded Mutant Mouse Resource & Research Center at the University of Missouri^49^. A successful transfer of GM_High_ was confirmed using 16S rRNA amplicon sequencing of cross-fostered pups and GM_High_-donating dams. GM_Low_ BTBR mice maintained their Jackson Laboratory-origin microbiome. Colonies of GM_Low_ and GM_High_ BTBR and B6 mice were then established. Mice used in the present study were from the 6^th^ to 8^th^ generation of their respective colonies.

Mice were group-housed under barrier conditions in microisolator cages (Thoren, Hazleton, PA, USA) on shaved aspen chip bedding with *ad libitum* access to autoclaved tap water and irradiated LabDiet 5053 chow (Labdiet, St. Louis, MO). Mice were maintained on a 12:12 light/dark cycle.

### BODY WEIGHTS

Neonatal body weights (D0 and D7) were collected using a tared New Classic MF #ML204 scale (Mettler Toledo; Columbus, OH, USA). Body weights at weaning (D21) and adulthood (D50) were measured using a Ranger^TM^ 3000 (OHAUS; Parsippany, NJ, USA).

### MICROBIOME ANALYSIS

#### SAMPLE COLLECTION

Fecal samples (1-2 pellets) were collected at necropsy from the distal colon of adult (D50) mice used in behavior testing and placed into 2 mL round-bottom tubes with a single 0.5 cm metal bead. Samples were flash frozen in liquid nitrogen then stored at −80°C until processing. Fecal DNA was extracted using a modified PowerFecal Pro Kit (QIAGEN; Hilden, North-Rhine-Westphalia, Germany). Briefly, lysis buffer (Solution C1) was directly added to the sample tube with the metal bead rather than the sample tube provided by the kit. Samples were then homogenized using a TissueLyser II (QIAGEN; Hilden, North-Rhine-Westphalia, Germany) for 10 min at 30 Hz before resuming extraction as prescribed by the manufacturer. DNA was eluted using Solution C6.

#### 16S rRNA TARGETED-AMPLICON SEQUENCING

Targeted-amplicon 16S rRNA library preparation and sequencing were performed by the University of Missouri Genomics Technology Core. Library preparations of the V4 region of the 16S rRNA gene were generated using PCR-amplification with the universal primers (U515F/806R)^50^ flanked by dual-index Illumina adapter sequences. PCR reactions each contained 100 ng metagenomic DNA, primers (0.2 µM each), dNTPs (200 µM each), and Phusion high-fidelity DNA polymerase (1U, Thermo Fisher, Waltham, MA, USA) in a 50 μL reaction. The amplification parameters were 98°C(3 min) + [98°C(15 s) + 50°C(30 s) + 72°C(30 s)] × 25 cycles + 72°C(7 min). Libraries were combined, mixed, and purified using Axygen Axyprep MagPCR clean-up beads for 15 min at room temperature. The products were washed multiple times with 80% ethanol and the dried pellet was resuspended in 32.5 µL of EB buffer (Qiagen, Venlo, The Netherlands), incubated for two minutes at room temperature, and then placed on a magnetic stand for five minutes. The amplicon pool was evaluated using an Advanced Analytical Fragment Analyzer automated electrophoresis system, quantified using quant-iT HS dsDNA reagent kits, and diluted according to the Illumina standard protocol for sequencing as 2 × 250 bp paired-end reads on the MiSeq instrument.

##### INFORMATICS

Sequences were processed using the Quantitative Insights into Molecular Ecology 2 v2021.8^51^. Paired-end reads were trimmed of the universal primers and Illumina adapters using *cutadapt*^52^. Reads were then denoised into unique amplicon sequence variants (ASVs) using DADA2^53^ with the following parameters: 1) reads were truncated to 150 bp in length, 2) reads with greater than 2 expected errors were discarded, 3) reads were merged with minimum overlap of 12 bp, and 4) chimeras were removed using the ‘consensus’ method. Unique sequences were filtered to between 249 and 257 bp in length. The remaining sequences were assigned a taxonomic classification using the *classify-sklearn* approach^54^ with the SILVA 138 99% NR reference database^55^ trimmed to the U515F/806R universal primers^50^.

The feature table of ASV counts per sample was rarefied to 28,850 ASVs per sample. The rarefied table was used for the remaining microbiome analyses. Chao1 and Shannon Indices (alpha diversity) were determined using the *microbiome*^56^ and *vegan*^57,58^ libraries, respectively. Beta diversity was compared by first creating a distance matrix with Bray-Curtis distances using the *vegan* library^57,58^. Differences in microbial beta diversity were visualized with principal coordinate analyses (PCoA) of quarter-root transformed feature tables with a Calliez correction using the *ape* library^59^. Differentially abundant taxa were identified using ALDEx2^26^ and ANCOM-BC2^27^ with a Benjamin-Hochberg^60^ corrected *p* value less than 0.05.

### BEHAVIORAL ASSAYS

All behavior tests were performed in a dedicated behavior suite separate from the animal housing room. Light levels for adult and neonatal testing were maintained at ∼5 *lux* and ∼100 *lux,* respectively. Sound levels were maintained at ∼45 dB during testing. Videos were captured using a DMK 22AUC03 IR camera (The Imaging Source; Charlotte, NC, USA) positioned 1.5 m above the cage bottom. Videos were recorded using ANY-maze v7.10 (ANY-maze; Wood Dale, IL, USA).

#### ULTRASONIC VOCALIZATION

USVs were collected using separation-induced vocalizations at PND7^21^. Briefly, cages were transferred to a behavior suite and allowed to acclimate for 60 min prior to testing. Neonates were individually separated from the dam and placed onto the floor of a heated, clean mouse cage enclosed within an isolated environmental chamber (Omnitech Electronics, Inc.; Columbus, OH, USA). An UltraSoundGate CM16 ultrasonic-sensitive microphone (AviSoft; Glienicke, Brandenburg, Germany) was suspended 15 cm above the cage bottom. The cage was then closed within the environmental chamber and USVs were recorded for a 5 min period using RECORDER USGH (AviSoft; Glienicke, Brandenburg, Germany). To prevent testing the same animal multiple times, mice were marked with a permanent marker after completing the recording, then returned to their birth dam. The recording chamber was cleaned with 70% EtOH before the first recording and after each subsequent trial. Mice were tested in alternating order of GM and sex as appropriate.

USV recordings were stored as *wav* files and analyzed using the machine-learning based *VocalMat* (v2021, <u>github.com/ahof1704/VocalMat</u>) using default settings^61^. VocalMat classifies individual mouse USVs into one of 12 classes: short, flat, chevron, reverse chevron, downward frequency modulation, upward frequency modulation, complex, multi steps, two steps, step down, step up, and noise. Calls classified as noise were removed from the vocalization rate and repertoire analysis. The vocal repertoire was determined by calculating the relative abundance of each call class for each mouse.

#### SELF-GROOMING TEST

Five-week-old mice were acclimated to an isolated behavior suite for 60 min prior to testing. Mice were individually placed into a clean, autoclaved cage and allowed to habituate for 10 min. Each mouse was then video recorded for the following 10 min^20^. A unique, randomly-generated identifier was placed within frame of each video, blinding the reviewer from strain, sex, and GM. Cages were cleaned prior to the first animal and after each trial using 70% EtOH. Each video was manually reviewed by a blinded reviewer. The total time spent grooming was measured using a stopwatch. Grooming behaviors included washing or scratching head, flank, limbs, and tail.

#### MARBLE BURYING TEST

Six-week-old mice were allowed to acclimate to an isolated behavior suite for 60 min prior to testing. Individual mice were placed in a standard Thoren mouse microisolator cage filled with 4-5 cm of aspen chip bedding and 12 black marbles placed in a 3 × 4 grid pattern on top of the bedding. Marbles were positioned prior to each trial using a template grid. Mice were placed into the cages and recorded for five minutes. A unique identifier was placed within frame of each video to blind the reviewer from strain, sex, and GM. Cages and marbles were cleaned prior to the first animal and after every trial using 70% EtOH. Fresh aspen chip bedding was provided for each trial. Videos were manually reviewed by a blinded reviewer using a stopwatch. Marble burying activity was defined as direct interaction with a marble or digging behavior.

#### SOCIAL PREFERENCE TEST

Seven-week-old mice were allowed to acclimate to an isolated behavior suite for 60 min prior to testing. Individual mice were placed into the center chamber of a three-chamber plexiglass arena (60 × 40 × 22 cm) and allowed to habituate to the arena for a 10 min period^28^. Following the acclimation period, the test mouse was enclosed in the middle chamber using plexiglass panels, blocking access to the outermost chambers. The test mouse was allowed to enter the middle chamber on its own volition without interaction from the experimenter. One cylindrical plexiglass cage (10 × 18.5 cm, 1 cm openings between vertical bars) was then placed in each of the outermost chambers in opposing corners. An age- and sex-matched A/J mouse (RRID:IMSR_JAX:000646, Jackson Laboratories; Bar Harbor, ME, USA) was placed in one of the cylinders (Stranger). A/J mice had been habituated to the cylinder during two 20 min training sessions, one on each side of the three-chamber arena. A plastic block was placed in the second cylinder (Object). The position of the stranger and object alternated between trials within each arena. The test mouse was then allowed to interact with the stranger or object for a period of 10 min. The position of the mouse and time spent in each zone was determined using ANY-maze. The three-chamber arena was cleaned with 70% EtOH prior to the first animal and after every subsequent test.

#### FOOD INTAKE

Food intake was monitored using a modified protocol from Cheathem et al.^13^. Food intake was monitored for four consecutive days for six consecutive weeks beginning after weaning (3 weeks old). On the first day (D0), the food hopper was topped off with LabDiet 5053 chow and the total hopper weight (hopper + chow) was measured using a OHAUS Ranger^TM^ 3000. Individual mouse weights were also recorded using the same scale. For the next three days, the hopper and mice were weighed in the same manner. The difference in hopper weights between each day was normalized to the combined animal weight of that cage. Feed efficiency was determined by normalizing the average food consumption to the combined cage weight.

Body and food intake weights were averaged for each week.

#### RUNNING WHEEL

Nine-week-old mice were housed in an animal room containing no other mice besides the animals undergoing wheel running evaluation. Mice were individually housed in Techniplast cages each containing a Low-Profile Wireless Running Wheel (Med Associates Inc.; Fairfax, VT, USA). Running wheels transmitted revolution counts to a central hub via Bluetooth. Locomotor activity was logged as revolutions per minute for one week using the Wheel Manager Data Acquisition Software from Med Associates. Total revolutions were converted to kilometers travelled using the prescribed conversion rate of (3.78 × 10^−4^ km/revolution).

### FECAL ENERGY LOSS

Fecal samples (799.1 ± 254.5 mg wet feces) were collected from seven-week-old mice over 2-3 morning collection periods. Fecal samples were dried at 65°C to a constant dry matter content. The gross energy (GE) of LabDiet 5053 chow and fecal samples was measured using a 6200 Isoperibol Calorimeter (Parr Instrument Co.; Moline, IL, USA). Benzoic acid (6318 ± 14 kcal GE/kg; Parr Instrument Co.) was used as the calibration standard. Temperature changes during combustion were monitored via a thermocouple, and the heat of combustion (ΔH) was calculated using the calorimeter’s specific heat capacity, then converted to caloric content (kcal/g feces). Accuracy corrections were applied to address background and ignition source heat.

### STATISTICAL ANALYSES

All statistical analyses were performed in R v 2021^62^. Differences in univariate data were assessed using two-way analysis of variance (ANOVA) with GM and sex as main effects. Longitudinal univariate data (i.e., food intake and physical activity) were assessed using three-way ANOVA tests with GM, sex, and time as main effects. *Post hoc* comparisons were made using a Tukey’s HSD test. Differences in paired data (i.e., social preference test) were assessed using paired T tests. All tests for significant differences in univariate data were performed using the *rstatix*^63^ package.

Differences in multivariate data (i.e., microbiome composition and vocal repertoire) were assessed using a permutational analysis of variance (PERMANOVA) using the *adonis2* function within the *vegan*^57,58^ library. Differentially abundant genera were identified using ALDEx2^26^ and ANCOM-BC2^27^. Significantly enriched taxa were identified by both tools as having a Benjamani-Hochberg-corrected *p* value < 0.05^64^.

## Acknowledgements

We would like to thank the Mutant Mouse Resource & Research Center at the University of Missouri (NIH U42 OD010918) for donating the surrogate GM_High_ CD-1 dams. ZM, KG, ALR, and ACE were supported by NIH U42 OD010918. ZM was also supported by NIH T32 GM008396.

## Data availability statement

All 16S rRNA sequencing data has been deposited to the National Center for Biotechnology Information (NCBI) Sequence Read Archive (SRA) under the BioProject number PRJNA1083497. All code has been deposited at https://github.com/ericsson-lab/btbr_2024.

## Supplementary Figure Legends

**Figure S1.**
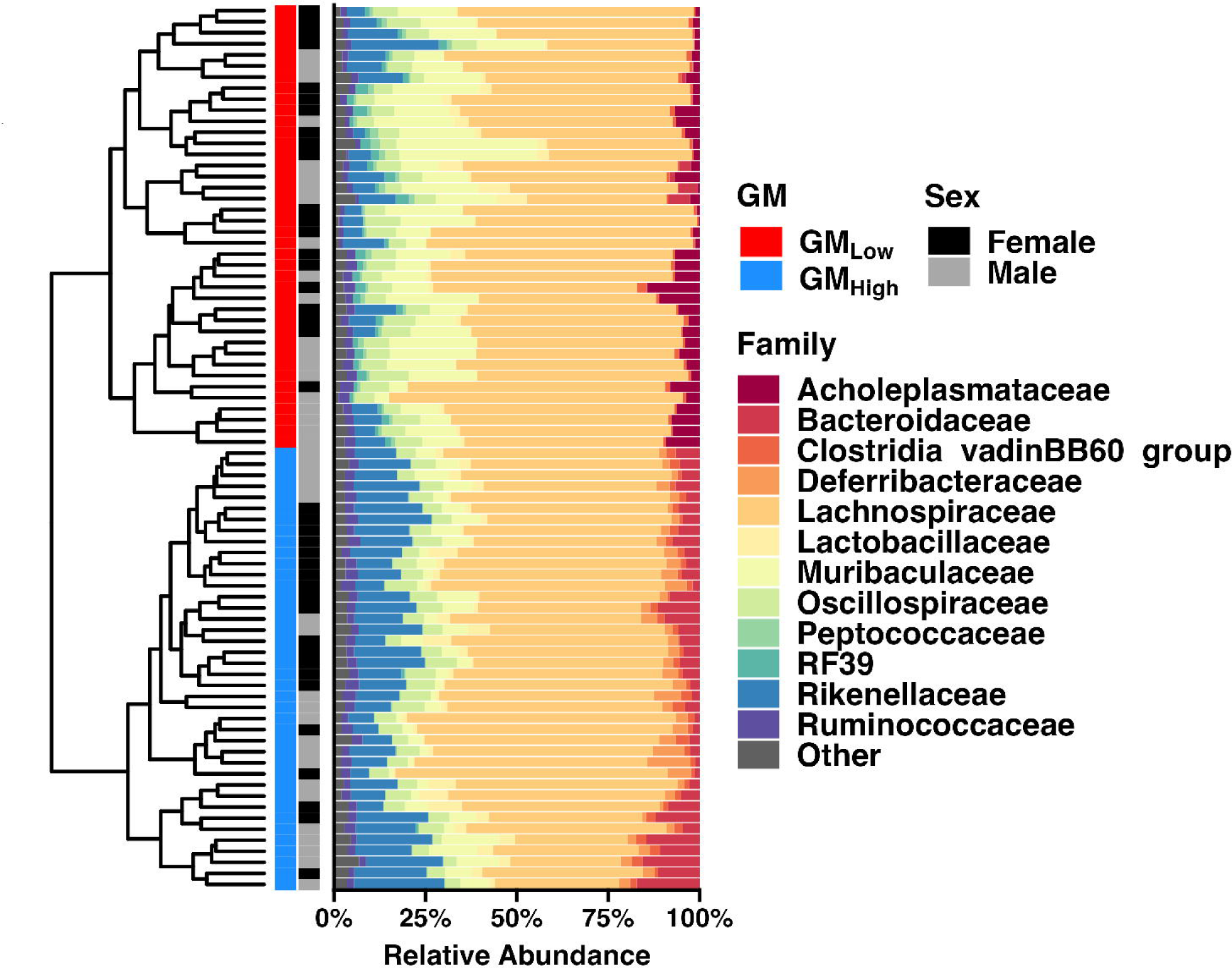
Standardized complex microbiomes differ in beta diversity and taxonomic composition. *Left:* Dendrogram depicting unsupervised hierarchical clustering of fecal microbiome samples based on composition. *Right*. Stacked bar chart depicting family-level abundance of dominant taxa (> 0.05%) in either GM.

**Figure S2.**
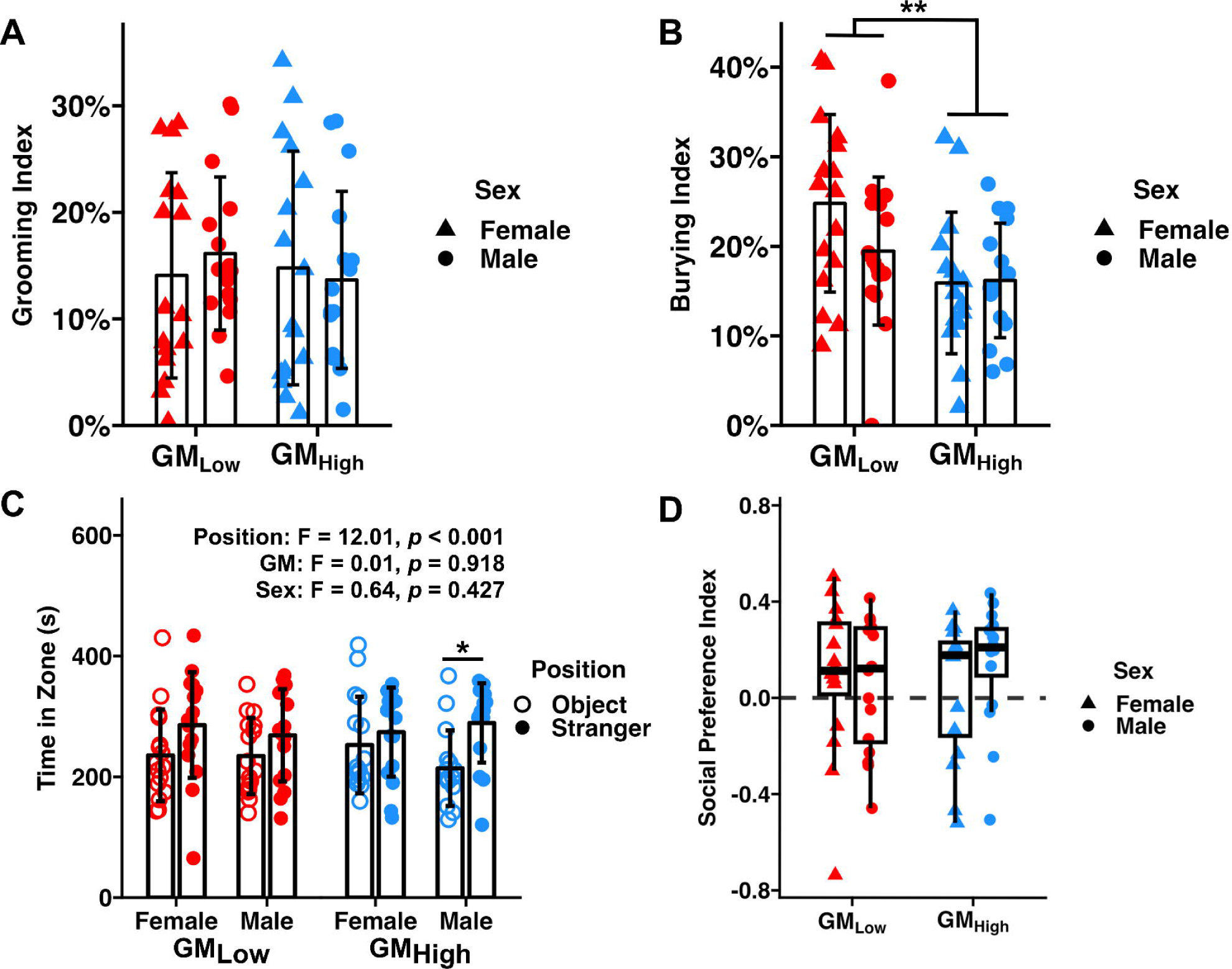
Standardized complex microbiomes selectively affect burying behavior of B6 mice. (A) Dot plot depicting Grooming Index. *p*_GM_ = 0.698, *p*_Sex_ = 0.838, Two-way ANOVA. (B) Dot plot depicting Burying Index. *\*\* p*_GM_ = 0.004, *p*_Sex_ = 0.224, Two-way ANOVA. (C) Dot plot depicting time spent in Stranger (closed circles) or Object (open circles) chambers of social preference test. * *p* = 0.027, Paired T test. Three-way ANOVA results are provided in inset. (D) Tukey box plot depicting Social Preference Index. *p*_GM_ = 0.771, *p*_Sex_ = 0.589, Two-way ANOVA.

**Figure S3.**
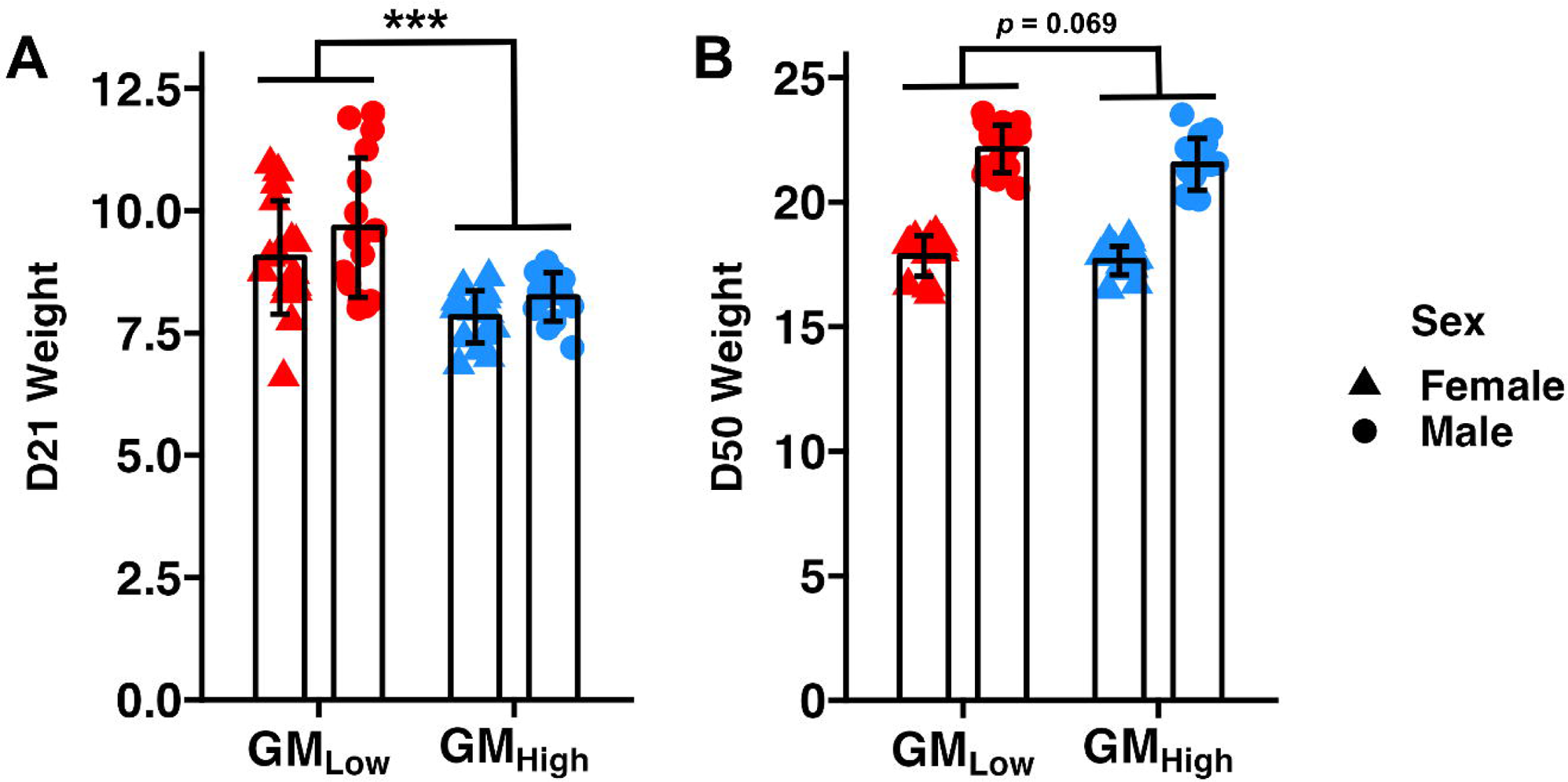
Standardized complex microbiomes affect body weight of B6 mice. Dot plots depicting body weights at D21 (A), and D50 (B) in GM_Low_ and GM_High_ B6 mice. *** *p*_GM_ < 0.001, Two-way ANOVA.

**Figure S4.**
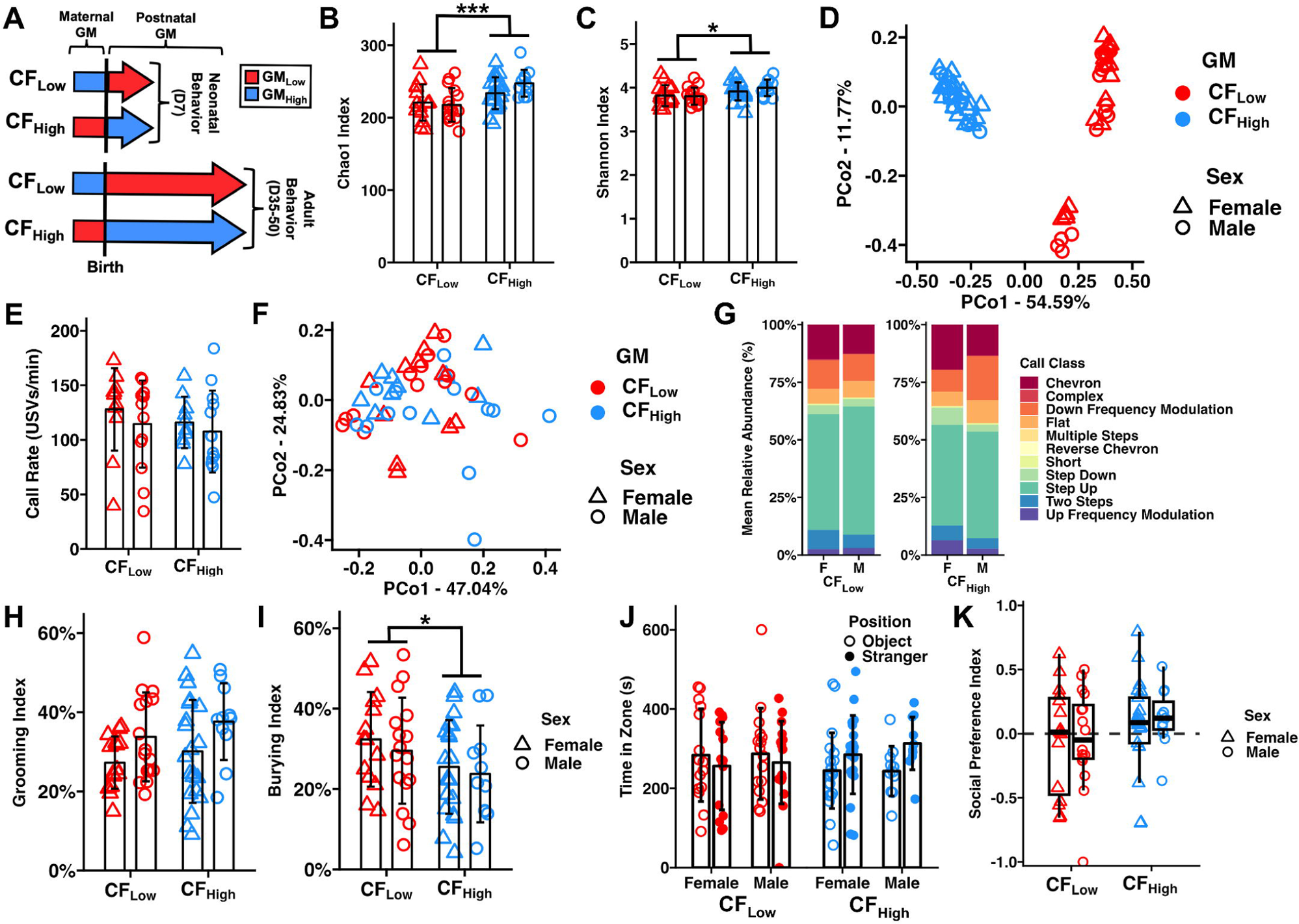
Cross-fostering abrogates select GM-mediated effects on male ASD-related behavior. (A) Graphical representation of cross-fostering experimental design depicting cohorts of neonatal (*n* = 10-13 mice/sex/GM) and adult (11-21 mice/sex/GM) BTBR mice. Inset depicts the GM that animals were exposed to pre- and postnatally. (B) Dot plot depicting Chao-1 Index. *** *p*_GM_ = 0.001, *p*_Sex_ = 0.401, Two-way ANOVA (C) Dot plot depicting Shannon Index. * *p*_GM_ = 0.013, *p*_Sex_ = 0.545, Two-way ANOVA (D) Principal coordinate analysis depicting Bray-Curtis dissimilarity between microbial communities. (E) Dot plot depicting USV rate. *p*_GM_ *=* 0.387, *p*_Sex_ = 0.306, Two-way ANOVA. (F) Principal coordinate analysis depicting Bray-Curtis dissimilarity of the relative abundance of USVs. *p*_GM_ *<* 0.001, *p*_Sex_ = 0.265, Two-way PERMANOVA. (G) Stacked bar charts depicting mean relative abundance of call types determined by VocalMat. (H) Dot plot depicting Grooming Index. *p*_GM_ *=* 0.237, *p*_Sex_ = 0.015, Two-way ANOVA. (I) Dot plot depicting Burying Index. * *p*_GM_ *=* 0.049, *p*_Sex_ = 0.474, Two-way ANOVA. (J) Dot plot depicting time spent in Stranger (closed circles) or Object (open circles) chambers of social preference test. (K) Tukey box plot depicting Social Preference Index. *p*_GM_ *=* 0.151, *p*_Sex_ = 0.741, Two-way ANOVA.

**Figure S5.**
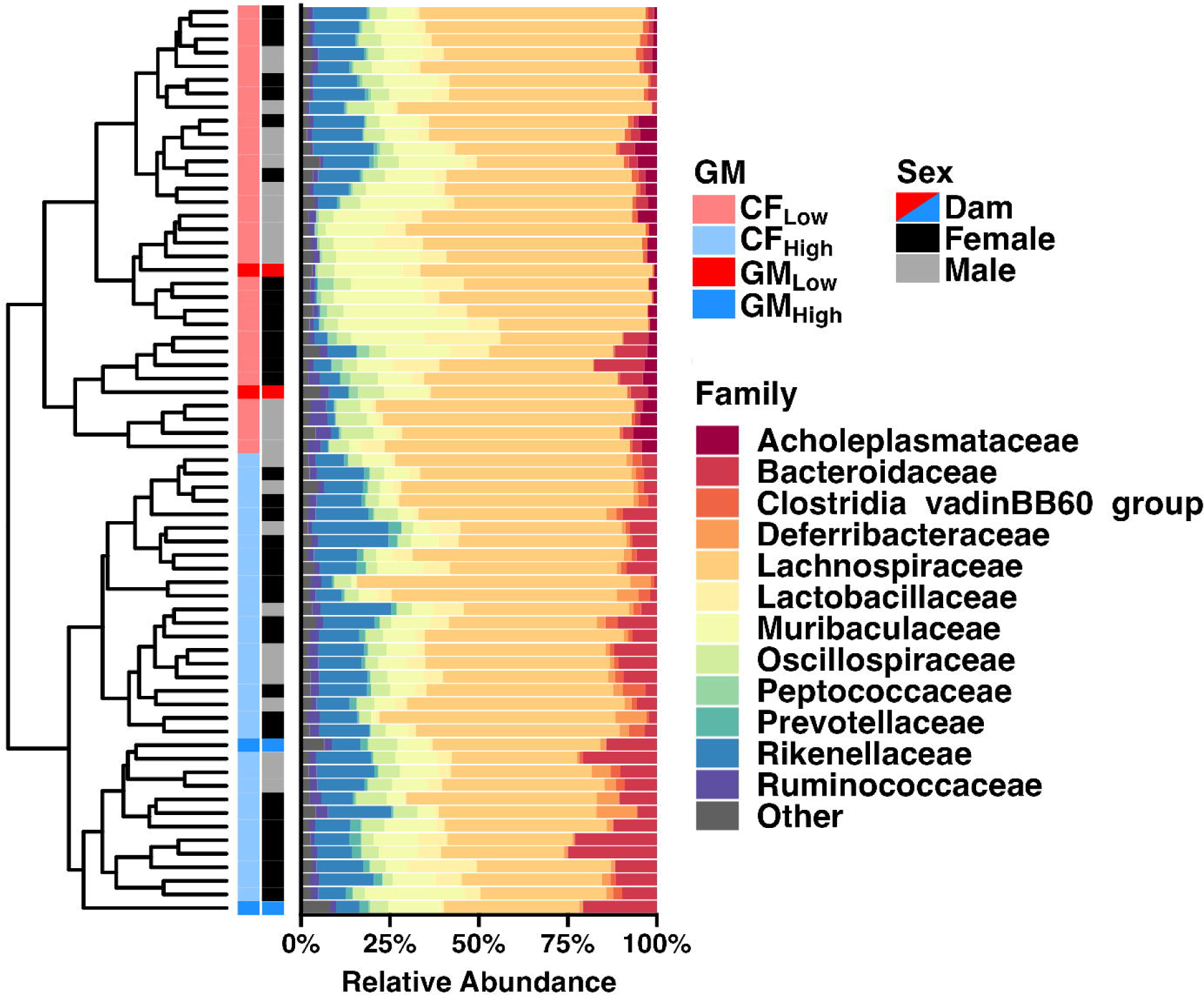
Standardized complex microbiomes were successfully transferred from surrogate dam to cross-fostered pup. *Left:* Dendrogram depicting unsupervised hierarchical clustering of fecal microbiome samples based on composition in relation to surrogate dam. *Right*. Stacked bar chart depicting family-level abundance of dominant taxa (> 0.05%) in either GM.

**Figure S6.**
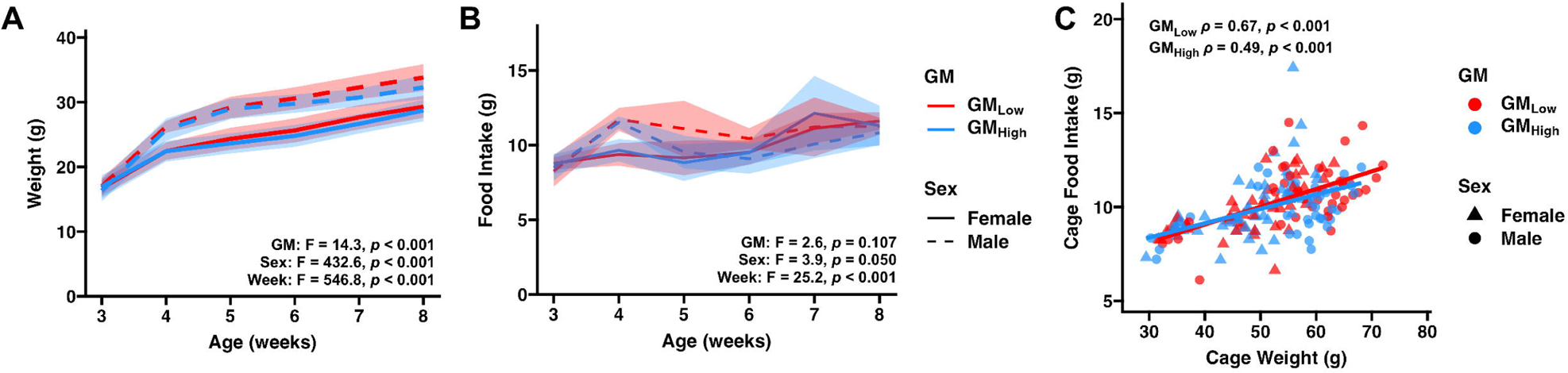
Standardized complex microbiomes affect body weight but not food intake. (A) Line plot depicting body weight observed in individual GM_Low_ and GM_High_ BTBR mice. Bold lines represent average body weight. Ribbon represents standard deviation. Inset depicts three-way ANOVA results. (B) Line plot depicting food intake of cages of pair-housed GM_Low_ and GM_High_ BTBR mice. Bold line represents average food intake. Ribbon represents standard deviation. Inset depicts three-way ANOVA results. (C) Dot plot depicting correlation between food intake of cage and body weight of mice within the same cage over the course of the experiment. Lines depict slope of correlation within either GM. Inset depicts correlation test results.

## References

1. Hirota T, King BH. Autism Spectrum Disorder. Jama 2023; 329:157–68.

2. Maenner MJ, Warren Z, Williams AR, Amoakohene E, Bakian AV, Bilder DA, Durkin MS, Fitzgerald RT, Furnier SM, Hughes MM, et al. Prevalence and Characteristics of Autism Spectrum Disorder Among Children Aged 8 Years — Autism and Developmental Disabilities Monitoring Network, 11 Sites, United States, 2020. Mmwr Surveill Summ 2023; 72:1–14.

3. Hung LY, Margolis KG. Autism spectrum disorders and the gastrointestinal tract: insights into mechanisms and clinical relevance. Nat Rev Gastroenterol Hepatol 2023; :1–22.

4. Morais LH, Schreiber HL, Mazmanian SK. The gut microbiota–brain axis in behaviour and brain disorders. Nat Rev Microbiol 2021; 19:241–55.

5. Bermudez-Martin P, Becker JAJ, Caramello N, Fernandez SP, Costa-Campos R, Canaguier J, Barbosa S, Martinez-Gili L, Myridakis A, Dumas M-E, et al. The microbial metabolite p-Cresol induces autistic-like behaviors in mice by remodeling the gut microbiota. Microbiome 2021; 9:157.

6. Cheng L, Wu H, Cai X, Zhang Y, Yu S, Hou Y, Yin Z, Yan Q, Wang Q, Sun T, et al. A Gpr35-tuned gut microbe-brain metabolic axis regulates depressive-like behavior. Cell Host Microbe 2024; 32:227–243.e6.

7. Desbonnet L, Clarke G, Shanahan F, Dinan TG, Cryan JF. Microbiota is essential for social development in the mouse. Mol Psychiatr 2014; 19:146–8.

8. Sgritta M, Dooling SW, Buffington SA, Momin EN, Francis MB, Britton RA, Costa-Mattioli M. Mechanisms Underlying Microbial-Mediated Changes in Social Behavior in Mouse Models of Autism Spectrum Disorder. Neuron 2019; 101:246–259.e6.

9. Kim S, Kim H, Yim YS, Ha S, Atarashi K, Tan TG, Longman RS, Honda K, Littman DR, Choi GB, et al. Maternal gut bacteria promote neurodevelopmental abnormalities in mouse offspring. Nature 2017; 549:528–32.

10. Sharon G, Cruz NJ, Kang D-W, Gandal MJ, Wang B, Kim Y-M, Zink EM, Casey CP, Taylor BC, Lane CJ, et al. Human Gut Microbiota from Autism Spectrum Disorder Promote Behavioral Symptoms in Mice. Cell 2019; 177:1600–1618.e17.

11. Ericsson AC, Davis JW, Spollen W, Bivens N, Givan S, Hagan CE, McIntosh M, Franklin CL. Effects of Vendor and Genetic Background on the Composition of the Fecal Microbiota of Inbred Mice. PLoS ONE [Internet] 2015; 10:e0116704. Available from: http://www.ncbi.nlm.nih.gov/pubmed/25675094

12. Ericsson AC, Hart ML, Kwan J, Lanoue L, Bower LR, Araiza R, Lloyd KCK, Franklin CL. Supplier-origin mouse microbiomes significantly influence locomotor and anxiety-related behavior, body morphology, and metabolism. Commun Biology 2021; 4:716.

13. Cheatham CN, Gustafson KL, McAdams ZL, Turner GM, Dorfmeyer RA, Ericsson AC. Standardized Complex Gut Microbiomes Influence Fetal Growth, Food Intake, and Adult Body Weight in Outbred Mice. Microorganisms 2023; 11:484.

14. Hart ML, Ericsson AC, Franklin CL. Differing Complex Microbiota Alter Disease Severity of the IL-10−/− Mouse Model of Inflammatory Bowel Disease. Front Microbiol 2017; 8:792.

15. Guo Y, Wang Q, Li D, Onyema OO, Mei Z, Manafi A, Banerjee A, Mahgoub B, Stoler MH, Barker TH, et al. Vendor-specific microbiome controls both acute and chronic murine lung allograft rejection by altering CD4+Foxp3+ regulatory T cell levels. Am J Transplant [Internet] 2019; 19:2705–18. Available from: https://www.ncbi.nlm.nih.gov/pubmed/31278849

16. Moskowitz JE, Doran AG, Lei Z, Busi SB, Hart ML, Franklin CL, Sumner LW, Keane TM, Amos-Landgraf JM. Integration of genomics, metagenomics, and metabolomics to identify interplay between susceptibility alleles and microbiota in adenoma initiation. BMC Cancer 2020; 20:600.

17. Kim E, Paik D, Ramirez RN, Biggs DG, Park Y, Kwon H-K, Choi GB, Huh JR. Maternal gut bacteria drive intestinal inflammation in offspring with neurodevelopmental disorders by altering the chromatin landscape of CD4+ T cells. Immunity 2021;

18. Choi GB, Yim YS, Wong H, Kim S, Kim H, Kim SV, Hoeffer CA, Littman DR, Huh JR. The maternal interleukin-17a pathway in mice promotes autism-like phenotypes in offspring. Science 2016; 351:933–9.

19. Moy SS, Nadler JJ, Young NB, Perez A, Holloway LP, Barbaro RP, Barbaro JR, Wilson LM, Threadgill DW, Lauder JM, et al. Mouse behavioral tasks relevant to autism: Phenotypes of 10 inbred strains. Behav Brain Res 2007; 176:4–20.

20. McFarlane HG, Kusek GK, Yang M, Phoenix JL, Bolivar VJ, Crawley JN. Autism-like behavioral phenotypes in BTBR T+tf/J mice. Genes Brain Behav 2008; 7:152–63.

21. Scattoni ML, Gandhy SU, Ricceri L, Crawley JN. Unusual Repertoire of Vocalizations in the BTBR T+tf/J Mouse Model of Autism. Plos One 2008; 3:e3067.

22. Moy SS, Nadler JJ, Young NB, Nonneman RJ, Segall SK, Andrade GM, Crawley JN, Magnuson TR. Social approach and repetitive behavior in eleven inbred mouse strains. Behav Brain Res 2008; 191:118–29.

23. Wöhr M, Roullet FI, Crawley JN. Reduced scent marking and ultrasonic vocalizations in the BTBR T+tf/J mouse model of autism. Genes Brain Behav 2011; 10:35–43.

24. Coretti L, Cristiano C, Florio E, Scala G, Lama A, Keller S, Cuomo M, Russo R, Pero R, Paciello O, et al. Sex-related alterations of gut microbiota composition in the BTBR mouse model of autism spectrum disorder. Sci Rep-uk 2017; 7:45356.

25. Golubeva AV, Joyce SA, Moloney G, Burokas A, Sherwin E, Arboleya S, Flynn I, Khochanskiy D, Moya-Pérez A, Peterson V, et al. Microbiota-related Changes in Bile Acid & Tryptophan Metabolism are Associated with Gastrointestinal Dysfunction in a Mouse Model of Autism. EBioMedicine 2017; 24:166–78.

26. Fernandes AD, Reid JN, Macklaim JM, McMurrough TA, Edgell DR, Gloor GB. Unifying the analysis of high-throughput sequencing datasets: characterizing RNA-seq, 16S rRNA gene sequencing and selective growth experiments by compositional data analysis. Microbiome 2014; 2:15–15.

27. Lin H, Peddada SD. Analysis of compositions of microbiomes with bias correction. Nature communications [Internet] 2020; 11:3514. Available from: https://www.ncbi.nlm.nih.gov/pubmed/32665548

28. Rein B, Ma K, Yan Z. A standardized social preference protocol for measuring social deficits in mouse models of autism. Nat Protoc 2020; 15:3464–77.

29. Daft JG, Ptacek T, Kumar R, Morrow C, Lorenz RG. Cross-fostering immediately after birth induces a permanent microbiota shift that is shaped by the nursing mother. Microbiome 2015; 3:17.

30. Yang M, Zhodzishsky V, Crawley JN. Social deficits in BTBR T + tf/J mice are unchanged by cross-fostering with C57BL/6J mothers. Int J Dev Neurosci 2007; 25:515–21.

31. Zhang Y, Gao D, Kluetzman K, Mendoza A, Bolivar VJ, Reilly A, Jolly JK, Lawrence DA. The maternal autoimmune environment affects the social behavior of offspring. J Neuroimmunol 2013; 258:51–60.

32. Caruso A, Ricceri L, Scattoni ML. Ultrasonic vocalizations as a fundamental tool for early and adult behavioral phenotyping of Autism Spectrum Disorder rodent models. Neurosci Biobehav Rev 2020; 116:31–43.

33. Peñagarikano O, Abrahams BS, Herman EI, Winden KD, Gdalyahu A, Dong H, Sonnenblick LI, Gruver R, Almajano J, Bragin A, et al. Absence of CNTNAP2 Leads to Epilepsy, Neuronal Migration Abnormalities, and Core Autism-Related Deficits. Cell 2011; 147:235–46.

34. Zhang Y, Li N, Li C, Zhang Z, Teng H, Wang Y, Zhao T, Shi L, Zhang K, Xia K, et al. Genetic evidence of gender difference in autism spectrum disorder supports the female-protective effect. Transl Psychiatry 2020; 10:4.

35. Willsey HR, Exner CRT, Xu Y, Everitt A, Sun N, Wang B, Dea J, Schmunk G, Zaltsman Y, Teerikorpi N, et al. Parallel in vivo analysis of large-effect autism genes implicates cortical neurogenesis and estrogen in risk and resilience. Neuron 2021; 109:788–804.e8.

36. Wang S, Wang B, Drury V, Drake S, Sun N, Alkhairo H, Arbelaez J, Duhn C, Genetics) TICG (TIC, Bromberg Y, et al. Rare X-linked variants carry predominantly male risk in autism, Tourette syndrome, and ADHD. Nat Commun 2023; 14:8077.

37. Vuong HE, Pronovost GN, Williams DW, Coley EJL, Siegler EL, Qiu A, Kazantsev M, Wilson CJ, Rendon T, Hsiao EY. The maternal microbiome modulates fetal neurodevelopment in mice. Nature 2020; 586:281–6.

38. Pronovost GN, Yu KB, Coley-O’Rourke EJL, Telang SS, Chen AS, Vuong HE, Williams DW, Chandra A, Rendon TK, Paramo J, et al. The maternal microbiome promotes placental development in mice. Sci Adv 2023; 9:eadk1887.

39. Flowers JB, Oler AT, Nadler ST, Choi Y, Schueler KL, Yandell BS, Kendziorski CM, Attie AD. Abdominal obesity in BTBR male mice is associated with peripheral but not hepatic insulin resistance. Am J Physiol-Endocrinol Metab 2007; 292:E936–45.

40. Queen NJ, Bates R, Huang W, Xiao R, Appana B, Cao L. Visceral adipose tissue-directed FGF21 gene therapy improves metabolic and immune health in BTBR mice. Mol Ther - Methods Clin Dev 2021; 20:409–22.

41. Takeuchi T, Kubota T, Nakanishi Y, Tsugawa H, Suda W, Kwon AT-J, Yazaki J, Ikeda K, Nemoto S, Mochizuki Y, et al. Gut microbial carbohydrate metabolism contributes to insulin resistance. Nature 2023; 621:389–95.

42. Zalewska A. Developmental milestones in neonatal and juvenile C57Bl/6 mouse – Indications for the design of juvenile toxicity studies. Reprod Toxicol 2019; 88:91–128.

43. Sanidad KZ, Zeng MY. Neonatal gut microbiome and immunity. Curr Opin Microbiol 2020; 56:30–7.

44. Russell AL, McAdams ZL, Donovan E, Seilhamer N, Siegrist M, Franklin CL, Ericsson AC. The contribution of maternal oral, vaginal, and gut microbiota to the developing offspring gut. Sci Rep 2023; 13:13660.

45. Oliphant K, Allen-Vercoe E. Macronutrient metabolism by the human gut microbiome: major fermentation by-products and their impact on host health. Microbiome 2019; 7:91.

46. Zeng X, Xing X, Gupta M, Keber FC, Lopez JG, Lee Y-CJ, Roichman A, Wang L, Neinast MD, Donia MS, et al. Gut bacterial nutrient preferences quantified in vivo. Cell 2022; 185:3441–3456.e19.

47. Xie C, Huang W, Young RL, Jones KL, Horowitz M, Rayner CK, Wu T. Role of Bile Acids in the Regulation of Food Intake, and Their Dysregulation in Metabolic Disease. Nutrients 2021; 13:1104.

48. Ruigrok RAAA, Weersma RK, Vila AV. The emerging role of the small intestinal microbiota in human health and disease. Gut Microbes 2023; 15:2201155.

49. Hart ML, Ericsson AC, Lloyd KCK, Grimsrud KN, Rogala AR, Godfrey VL, Nielsen JN, Franklin CL. Development of outbred CD1 mouse colonies with distinct standardized gut microbiota profiles for use in complex microbiota targeted studies. Sci Rep 2018; 8:10107.

50. Caporaso JG, Lauber CL, Walters WA, Berg-Lyons D, Lozupone CA, Turnbaugh PJ, Fierer N, Knight R. Global patterns of 16S rRNA diversity at a depth of millions of sequences per sample. Proc Natl Acad Sci [Internet] 2011; 108:4516–22. Available from: http://www.ncbi.nlm.nih.gov/pubmed/20534432

51. Bolyen E, Rideout JR, Dillon MR, Bokulich NA, Abnet CC, Al-Ghalith GA, Alexander H, Alm EJ, Arumugam M, Asnicar F, et al. Reproducible, interactive, scalable and extensible microbiome data science using QIIME 2. Nat Biotechnol [Internet] 2019; 37:852–7. Available from: https://www.ncbi.nlm.nih.gov/pubmed/31341288

52. Martin M. Cutadapt removes adapter sequences from high-throughput sequencing reads. EMBnetJ [Internet] 2011; 17:10–2. Available from: https://journal.embnet.org/index.php/embnetjournal/article/view/200

53. Callahan BJ, McMurdie PJ, Rosen MJ, Han AW, Johnson AJ, Holmes SP. DADA2: High-resolution sample inference from Illumina amplicon data. Nature methods [Internet] 2016; 13:581–3. Available from: https://www.ncbi.nlm.nih.gov/pubmed/27214047

54. Kaehler BD, Bokulich NA, McDonald D, Knight R, Caporaso JG, Huttley GA. Species abundance information improves sequence taxonomy classification accuracy. Nat Commun 2019; 10:4643.

55. Pruesse E, Quast C, Knittel K, Fuchs BM, Ludwig W, Peplies J, Glockner FO. SILVA: a comprehensive online resource for quality checked and aligned ribosomal RNA sequence data compatible with ARB. Nucleic Acids Res [Internet] 2007; 35:7188–96. Available from: https://www.ncbi.nlm.nih.gov/pubmed/17947321

56. Lahti L, Shetty S. microbiome R package. 2012; Available from: https://microbiome.github.io/microbiome/

57. Dixon P. VEGAN, a package of R functions for community ecology. J Veg Sci 2003; 14:927–30.

58. Oksanen J, Blanchet FG, Kindt R, Legendre P, Minchin PR, O’Hara RB, Simpson GL, Solymos P, Stevens MHH, Wagner H. Vegan: Community Ecoloy Package. R package version 2.2-0. 2014; Available from: https://github.com/vegandevs/vegan

59. Paradis E, Schliep K. ape 5.0: an environment for modern phylogenetics and evolutionary analyses in R. Bioinformatics [Internet] 2019; 35:526–8. Available from: https://academic.oup.com/bioinformatics/article/35/3/526/5055127

60. Y Hyb. Controlling the false discovery rate: a practical and powerful approach to multiple testing. Journal of the Royal Statistical Society Series B 1995; 57:289–300.

61. Fonseca AH, Santana GM, Ortiz GMB, Bampi S, Dietrich MO. Analysis of ultrasonic vocalizations from mice using computer vision and machine learning. eLife 2021; 10:e59161.

62. Team RDC. R: A Language and Environment for Statistical Computing. 2022; Available from: http://www.R-project.org

63. Kassambara A. rstatix: Pipe-Friendly Framework for Basic Statistical Tests. 2022; Available from: https://CRAN.R-project.org/package=rstatix

64. Benjamini Y, Hochberg Y. Controlling the False Discovery Rate: A Practical and Powerful Approach to Multiple Testing. J Royal Statistical Soc Ser B Methodol [Internet] 1995; 57:289–300. Available from: https://rss.onlinelibrary.wiley.com/doi/10.1111/j.2517-6161.1995.tb02031.x

